# Nanovibrational stimulation preferentially enhances osteogenic responses in zebrafish

**DOI:** 10.64898/2026.06.18.733105

**Authors:** Opeyemi Adigun, Mengdi Wang, Jonathan A. Williams, Jérémie Zappia, Oana Dobre, Merna Maung, Stuart Reid, Peter G. Childs, Joanna J. Moss, Chrissy L. Hammond

**Affiliations:** School of Biochemistry and Biomedical Sciences, University of Bristol, Bristol, BS8 1TD, UK; Centre for the Cellular Microenvironment, Department of Biomedical Engineering, University of Strathclyde, Glasgow, G1 1XQ, UK; Centre for the Cellular Microenvironment, Advanced Research Centre, University of Glasgow, Glasgow, G11 6EW, UK

## Abstract

Mechanical cues are key regulators of bone formation, yet their potential as therapeutic stimuli remains incompletely explored in vivo. Nanovibrational stimulation, which delivers low-amplitude, high-frequency mechanical input, has been shown to promote osteogenic differentiation in vitro, and rodent studies have similarly demonstrated osteogenic effects. However, its impact on other cellular systems at the whole-organism level remains poorly understood. Here, we demonstrate that nanovibrational stimulation enhances myogenic differentiation in vitro before investigating its effects on skeletal development and tissue specificity in zebrafish.

Larval zebrafish exposed to nanovibrational stimulation exhibited increased osteoblast numbers and enhanced bone formation relative to controls. In adult zebrafish, nanovibration increased the osteoblast response to a fracture-like injury, indicating enhanced osteogenic activity during repair. To assess tissue specificity, we examined additional cell types and systems relevant to skeletal regeneration, including cartilage, muscle, vasculature and innate immune cells. Although nanovibrational stimulation promoted myogenic differentiation of C2C12 cells in vitro, its effects on muscle and other non-skeletal tissues in zebrafish larvae were comparatively limited. These findings suggest that nanovibrational stimulation exerts a preferential effect on osteogenic processes in vivo.

These findings demonstrate that nanovibrational stimulation preferentially enhances osteoblast-mediated bone formation and skeletal injury responses in zebrafish, without causing significant perturbations in other tissues. Our results establish zebrafish as a tractable in vivo model for investigating vibration-induced mechanobiological processes, providing a more physiologically relevant representation of tissue-level mechanotransduction than conventional two-dimensional culture systems. Furthermore, these findings highlight the potential of nanovibrational stimulation as a non-invasive strategy to promote bone regeneration and fracture repair.

## Introduction

The restoration of bone following fracture remains a significant clinical challenge, particularly in cases of delayed healing or non-union. While current therapeutic strategies largely focus on biochemical and surgical interventions, there is increasing interest in the role of mechanical cues as regulators of skeletal repair ^1,2^ Bone is a highly mechanosensitive tissue, and physical stimuli, ranging from macroscopic loading to microscale substrate properties, are known to influence osteogenesis and fracture repair across vertebrate species ^13,4^. More recently, vibrational cues have emerged as a distinct class of biophysical stimulus, with nanoamplitude vibrational stimulation offering a potential route to direct osteogenic cell fate and enhance bone formation^5,6^,.

Vibrational stimulation is thought to enhance myogenic differentiation by activating similar mechanosensitive pathways within the cell to those activating osteogenesis^8–11^. Mechanical cues at this scale are sensed by the cytoskeleton and transmitted to the nucleus, where they promote the expression of myogenic transcription factors, including MyoD and myogenin, that drive commitment towards a muscle phenotype and the subsequent fusion of myoblasts into elongated, multinucleated myotubes. This mirrors the role of physical forces in native skeletal muscle development, where mechanical loading from surrounding tissue is a key regulator of satellite cell activation and myoblast fusion. Vibration may also increase cytoskeletal tension and focal adhesion assembly, further reinforcing the pro-differentiation signaling cascade and facilitating the cell–cell contacts required for myotube elongation.

Several vibration-based mechanical stimulation strategies have been investigated for bone repair and regeneration, including low-magnitude mechanical signals, whole-body vibration, low-intensity pulsed ultrasound, and shockwave therapy. These interventions vary widely in their mode of action, ranging from whole-limb or muscle-mediated skeletal loading to externally applied pressure waves^8,12–14^. Nanovibrational stimulation is distinct from these approaches in that it applies extremely low-amplitude, high-frequency oscillatory cues directly to cells or tissues, typically at 1 kHz with nanoscale displacement amplitudes of approximately 30 nm. These parameters allow the application of nanonewton-level forces at the cellular level, sufficient to activate certain cellular mechanosensors^15,16^ .

A growing body of work demonstrates that such stimulation can drive osteogenic differentiation of mesenchymal stem cells (MSCs) *in vitro*, increasing the expression of osteogenic markers, including osteopontin and osteocalcin, and promoting matrix mineralisation even in the absence of exogenous osteoinductive factors^5,1,18^. Recent studies have further shown that nanovibration induces coordinated metabolic and lipidomic changes associated with osteodifferentiation, including shifts in energy demand, membrane remodelling, and redox regulation^19^. Mechanistically, these effects have been linked to activation of mechanotransductive pathways involving cytoskeletal remodelling (e.g. ERK1/2 and Wnt/β-catenin pathways), ion channel signalling (e.g. Piezo1 and TRPV1), and downstream transcriptional regulators of osteogenesis. In addition, nanovibrational stimulation has been shown to modulate inflammatory and reactive oxygen species (ROS)-related signalling, which may play a permissive role in osteogenic commitment in three-dimensional culture systems^20^.

Importantly, recent advances in nanovibrational bioreactor systems have enabled reproducible induction of osteoblast-like phenotypes across both two- and three-dimensional culture environments, supporting the translational potential of this approach However, despite these advances in cell culture models, much of this work has focused on bone-related cell types cultured in isolation, limiting understanding of how other musculoskeletal tissues respond to comparable vibrational inputs. In addition, conventional monolayer culture systems provide a limited representation of the spatial, structural, and mechanical complexity of the native ECM found in musculoskeletal tissues. This is particularly important given that matrix composition, dimensionality, and cell-matrix coupling are likely to influence force transmission and mechanosensitive signaling as are interactions between tissues and that these will change through ontogeny. The extent to which nanovibrational stimulation can selectively influence the cells of the skeletal tissues *in vivo*, without broadly perturbing other cell types and organ systems, remains incompletely understood. In previous work, a wearable nanovibration delivery system was developed and applied in a rat model of spinal cord injury-induced osteoporosis, transmitting a 1 kHz, 30 nm nanoscale stimulus to the paralysed hindlimbs. Although the intervention did not reverse the severe established osteoporosis present in this model, it was associated with increased circulating levels of the bone formation marker procollagen type I N-terminal propeptide (P1NP), consistent with a biological effect on bone formation. However, the cellular and tissue-level specificity of nanovibrational stimulation in vivo remains unclear. The zebrafish (*Danio rerio*) provides a powerful vertebrate model for addressing cellular effects of nanovibration *in vivo*. The optical translucency of larvae during early development, genetic tractability, and conserved skeletal development pathways make them well-suited for studying osteogenesis and mechanobiology *in vivo*^21–25^. Furthermore, adult zebrafish scales constitute a tractable and physiologically relevant model of adult bone remodelling post injury, containing functional osteoblasts and osteoclasts as well as a mineralised matrix ^26–29^.

In this study, we investigate the effects of nanovibrational stimulation (1 kHz, 30 nm) on bone in zebrafish skeletal tissues. We quantify osteoblast number and bone formation during larval development, and osteoblast recruitment to a skeletal injury in adult elasmoid scales. To assess the specificity of nanovibrational effects, we also examine a range of additional tissues and cell types pertinent to skeletal remodelling, including craniofacial cartilage, innate immune cells, and skeletal muscle fibres in both trunk and craniofacial regions.

## Materials and Methods

Nanovibrational stimulation was delivered using a previously described nanovibrational bioreactor platform (Figure 1A)^15^. The system consists of an array of 13 piezoelectric actuators topped by a magnetic sample receiving plate, allowing cultureware to be firmly bonded to the vibrating surface by self-adhesive magnetic sheet. During operation, the actuator array drives the top plate in a nominally uniform, pistonic vertical motion, so that the cultureware is displaced sinusoidally at 1 kHz with an intended nanoscale amplitude of approximately 30 nm.

**Figure 1.**
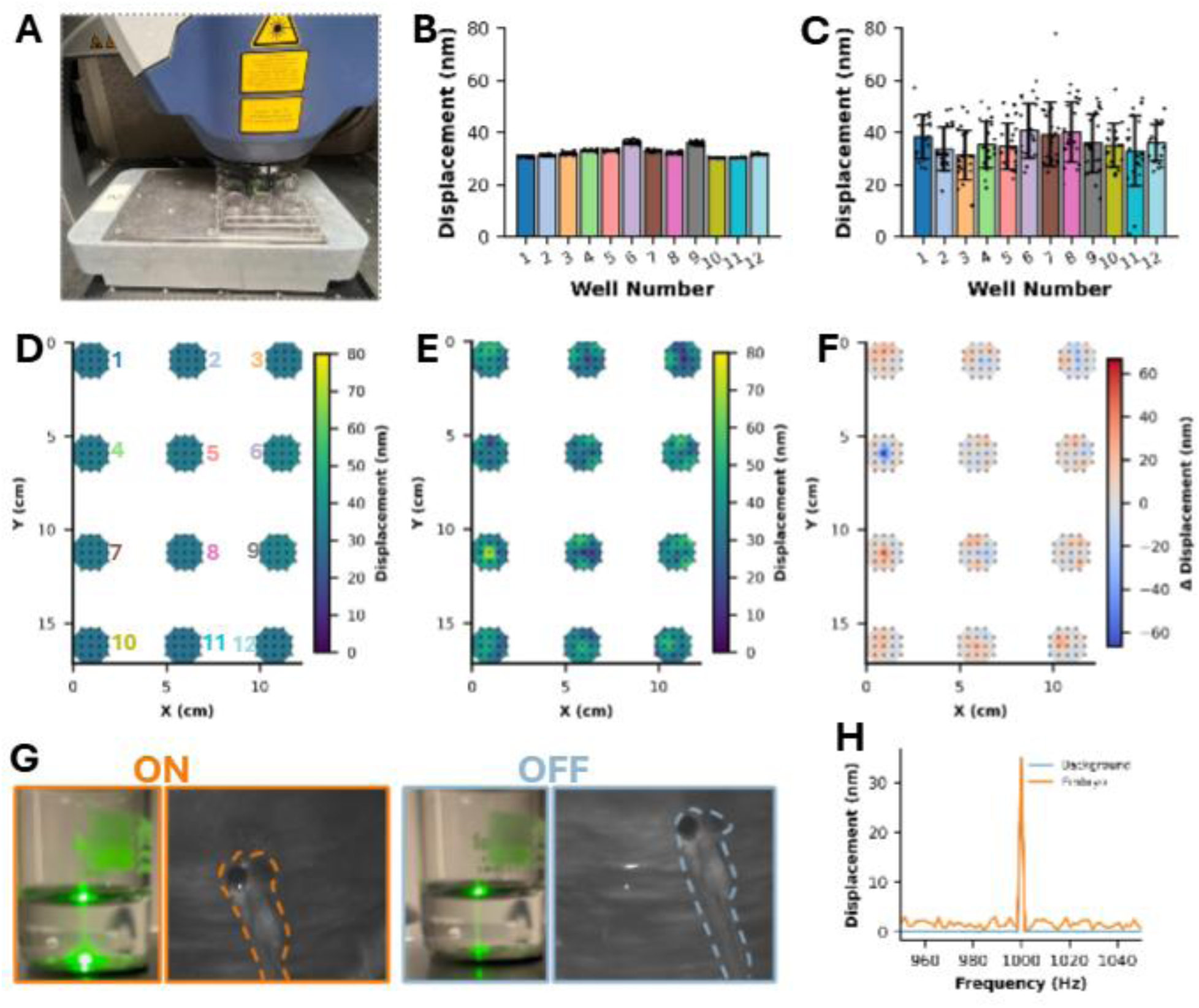
Laser Doppler vibrometry characterisation of nanovibrational stimulation in zebrafish culture systems. **(A)**Experimental setup showing the scanning laser Doppler vibrometer(LDV) used to characterise vibration transmitted from the nanovibrational bioreactor to cultureware. The bioreactor was operated at 1 kHz with a nominal displacement amplitude of 30nm, delivering a vertical pistonic stimulus to magnetically 12-wellplates. **(B)** Maximum in-phase displacement measured at the bottom surface of each well of the empty 12-well plate. **(C)**Maximum in-phase displacement measured at the water surface in each well following the addition of 2 mL of water. **(D)** Spatial heatmap representation of the bottom-surface displacement measurements. **(E)** Spatial heat map representation of the water-surface displacement measurements. **(F)** Difference displacement map showing the difference between the displacement measured at the well bottom and at the water surface. **(G)** Experimentalsetup used for embryo-targeted LDV measurements, showing laser positioning on a zebrafish embryo and on an adjacent background region. A measurable laser signal was obtained in both cases. **(H)** Representative frequency-domain displacement spectra acquired with the LDV focused on a zebrafish embryo and on adjacent background regions

### Validation of the nanovibration setup in multiwell format

The vibrational response of the experimental zebrafish culture setup was characterised using a scanning laser Doppler vibrometer (LDV) (Model MSA-100-3D, Polytec GmbH, Germany). The setup comprised a standard 12-well plate used for zebrafish embryo culture, magnetically coupled to the top plate of the nanovibrational bioreactor using a 2 mm-thick adhesive rubber magnet sheet cut to size. The bioreactor was operated in its standard mode, delivering a nominal vertical pistonic sinusoidal vibration at 1 kHz with a displacement amplitude of 30 nm to the coupled cultureware. For LDV measurements, the excitation signal was generated using the LDV’s internal signal generator and amplified using an external Behringer KM750 amplifier, operated at maximum gain, to drive the piezo array. The excitation signal was a 1 kHz sine wave with a generator amplitude of 130 mV and zero offset, corresponding to a nominal external amplification of approximately 40×. The LDV was operated in FFT acquisition mode, with measurements acquired in the vertical (+Z) direction using velocity output. Acquisition settings were: 12.5 kHz sampling frequency, 5 kHz bandwidth, 3200 FFT lines, 640 ms sample time, 1.5625 Hz frequency resolution, rectangular window, no filtering, and complex averaging over 10 acquisitions. In-phase displacement was measured in each of the 12 wells of the culture plate. For each well, a grid of 21 evenly spaced measurement points, covering a central area of approximately 3.0 cm², centred on the well’s midpoint, was defined in the PSV software (Polytec GmbH, Germany). Measurements were first acquired from the bottom surface of the wells in the absence of water to characterise vibration transmission through the plate itself. The wells were then filled with 2 mL of water, and the same 21 measurement positions were used to acquire measurements from the water surface. Using identical point locations for the well bottom and water surface enabled direct comparison of displacement at corresponding positions within each well.

To examine whether the nanovibrational signal could be resolved from individual zebrafish embryos, separate LDV measurements were performed in a small 10 mL glass beaker (Figure 1G) rather than in the 12-well plate, as optical visualisation within the wells was insufficient for reliable laser targeting. The glass vessel provided improved optical access and enabled the laser to be focused either directly on an embryo or on an adjacent background region at the same depth. Zebrafish embryos naturally resided in the lower region of the vessel, close to the vibrating bottom surface, and representative spectra and peak displacement amplitudes at 1 kHz were acquired from both embryo-targeted and background-targeted positions. These measurements were used to assess whether any distinct embryo-associated signal could be resolved relative to the surrounding vibrating environment.

### Statistics for nanovibration analysis

Statistical analysis was performed in Python using the pandas, NumPy, and SciPy libraries. Displacement data were assigned to wells following spatial clustering of measurement coordinates. For each well, summary statistics were calculated and are presented as the mean ± standard deviation. Pairwise differences between wells were assessed using two-sided Welch’s t-tests assuming unequal variances, with Holm–Bonferroni correction applied to control for multiple comparisons.

**C2C12 cell culture and nanovibration**_C2C12 myoblasts were seeded in 12-well plates at 20,000 cells per well and cultured in standard growth medium until approximately 80% confluent. Cells were then allocated to control or nanovibration conditions and maintained either in growth medium or in differentiation medium. Differentiation was induced by replacing the growth medium with DMEM supplemented with 1% penicillin/streptomycin, 0.1% FBS, and 1% insulin-transferrin-selenium (ITS), at this point Nanovibration was applied continuously for 48 h at 1 kHz and approximately 30 nm nominal displacement amplitude, in differentiation medium prior to fixation and immunostaining.

**C2C12 immunostaining**_Cells were fixed in 4% paraformaldehyde and permeabilised with 0.1% Triton X-100 for 10 min. Samples were blocked with 2% BSA in PBS for 45 min at room temperature, followed by overnight incubation at 4 °C with an Alexa Fluor 488-conjugated MF20 monoclonal antibody against myosin heavy chain/Myosin 4 (eBioscience™, Product #53-6503-82), diluted 1:200 in 0.1% /PBS. Cells were washed with 0.5% Tween-20 and counterstained with DAPI for 10 min at room temperature before confocal imaging.

**Imaging and quantification of myotube formation**_Immunofluorescence images were acquired using a Zeiss LSM 900 confocal microscope. MF20 staining was used to identify differentiated myotubes, and DAPI staining was used to identify nuclei.

### Zebrafish husbandry

Zebrafish (Danio rerio) were raised and maintained under standard conditions. Experiments were approved by the local ethics committee (the Animal Welfare and Ethical Review Body of the University of Bristol) and performed under UK Home Office project license. Transgenic lines used: *Tg(Ola. Sp7:nlsGFP)*^zf132^ for osteoblasts^30^, *TgBAC(col2a1a:mCherry*)*^hu^*^5900^ for chondrocytes^31^, *Tg(smyhc1:EGFP)^i^*^104^ for muscle^32^, *Tg(mpeg1:mCherry)* for macrophages^33^ and *TgBAC(mpx:GFP)* for neutrophils^34^.

### Zebrafish anesthesia

Recovery anesthesia was performed by diluting a stock solution of Tricaine Methanesulphonate (MS222) into Danieau solution for larvae (100 mg/L). Larvae were recovered in a fresh Danieau solution. For non-recovery anesthesia, a stock solution of Tricaine Methanesulphonate (MS222) was dissolved in Danieau solution for larvae (640 mg/L).

### Whole-mount zebrafish immunohistochemistry

Larvae at 5 days post-fertilisation (dpf) were fixed in 4% PFA then stored at -20°C in 100% MeOH. Larvae were re-hydrated, washed in PBS-T (PBS with 0.1% Triton X-100) before permeabilisation with 15 µg/mL proteinase K (4333793, Sigma) at 37 °C for 1 hour. For A4.1025 antibody staining, larvae were instead permeabilised in 0.25% Trypsin for 15 minutes on ice. Next, samples were blocked in 5% horse serum and incubated with primary antibodies overnight at 4 °C (anti-myosin (clone A4.1025) (sc-53088, 1:500, Insight Biotech, UK); anti-col2a1 (II-II6B3-s, 1:20, DSHB, IA, USA)). Samples were washed extensively in 1X PBS-T before incubation with Alexa-Fluor secondary antibodies (Invitrogen) diluted at 1:500 in 5% horse serum. Incubations with secondary antibody were performed for 2 hours at room temperature, in the dark, then imaged on a confocal microscope.

### Zebrafish cell proliferation assay

Cell proliferation was measured using the Click-iT EdU imaging kit (Invitrogen, C10337) according to the manufacturer’s instructions. Briefly, 5 dpf larvae were treated with 400 μM EdU in Danieau’s buffer for 24 hours. Larvae were fixed in 4% PFA overnight at 4 °C, washed, and then incubated in the Click-iT reaction cocktail for 30 minutes, following the manufacturer’s instructions.

### Live bone stains

Live larval or adult fish were incubated for 1 hour in 0.1 % Alizarin Red S (Sigma, A5533) in Danieau solution. Adult scales were incubated in 40 μM calcein powder (pH 8.0) (Sigma-Aldrich, C0875) in Danieau solution for 30 minutes. For both, fish were rinsed twice and imaged on a stereomicroscope.

### Stereomicroscope imaging

Live larval zebrafish or adult scales were imaged using a DFC700T camera mounted to a Leica MZ10F modular stereomicroscope system (Leica Microsystems) at 1-12× magnification. For live imaging, fish were anaesthetised using 0.1 mg/mL MS222 (Tricaine methane sulfonate) diluted in Danieaus and imaged laterally.

### Confocal microscope imaging

Larvae were mounted in 1% LMP agarose (16520050, Thermofisher, MA, USA) and imaged using a Leica SP5-II AOBS tandem scanner confocal microscope attached to a Leica DMI 6000 inverted epifluorescence microscope with oil immersion 20× or 40× objectives run using Leica LAS AF software (Leica, Germany). Maximum intensity projection images were assembled using LAS AF Lite (Leica) and Fiji ^35^.

### Elasmoid scale culture

Scales from anaesthetised adult zebrafish carrying the osterix transgene *Tg(Ola. Sp7:nlsGFP)*^zf132^ were plucked from the lateral midline near the dorsal fin using forceps and 5-8 scales were transferred to 6 well plates with 2 mL osteogenic scale culture media (o-DMEM) per well (Dulbecco’s Modified Eagle Medium without phenol red (DMEM; #31053, Gibco Life Technologies), 2% penicillin/streptomycin (#15140, Gibco Life Technologies), 1% FBS (HyClone #SH30072.03), 1% GlutaMAX 100X (Gibco Life Technologies), 1 mM Sodium pyruvate (#S8638, Sigma), 4 mM calcium chloride, 10 mM beta-glycerophosphate) as previously described^26^. Plates were incubated at 28 °C with 5% CO_2_ for 48 hours, with one plate on the nanovibration platform and the control plate on an adjacent shelf.

### Scale bone injury model

In a 5 cm petri dish containing o-DMEM, a sterile scalpel was used to create a single fracture on the lower lateral region of each scale, extending from the circulus towards the focus. Scales were transferred into multi-well plates and cultured for 48 hours, with or without nanovibration treatment. Scales were imaged on the Evident VS200 slide scanner with a 20× objective. Following imaging, mean pixel intensity at the injury site was quantified using a fixed sized region of interest (ROI) applied consistently across all scales. To account for background fluorescence and inter-scale variability, the mean pixel intensity from the uninjured side of each scale was measured and subtracted from the fracture site mean pixel intensity. The resulting values were used as the normalised mean pixel intensity for the subsequent analysis.

### Image analysis and statistics

All image analysis was performed using Image J (Fiji)^35^. Operculum measurement, jaw cartilage and muscle, and somite measurements were made manually in Fiji using the straight line or polygon tool from max intensity projections of confocal z-stacks. For EdU cell proliferation counts, EdU-positive muscle cells positive for A4.1025 were counted by going through z-stacks. EdU positive cells in or overlying the somite that were not positive for A4.1025 were not included.

Statistical analyses were performed using GraphPad Prism v.11.0.0. Error bars on all graphs represent the mean ± standard deviation. Statistical tests and specific parameters for each assay are mentioned in the appropriate figure legend.

## Results

### Nanovibration passes through larvae and scales in multiwell culture plates

Scanning laser Doppler vibrometry was used to characterise vibration transmission through the 12-well plate used for zebrafish embryo and scale culture experiments (Figure 1A). In-phase maximum displacement measurements acquired from the bottom surface of the empty 12-well plate confirmed delivery of consistent nanoscale vibration across the culture wells (Figure 1B and D). Mean well displacements ranged from 30 to 36 nm, with low within-well variability, indicating stable vibration across the measured area of each well.

In comparison with the well-bottom measurements, displacement recorded at the water surface showed greater variability, both within and between wells, as expected for a freely moving liquid interface (Figure 1C, E and F). Nevertheless, all wells exhibited clear nanoscale displacement at the water surface, with mean values ranging from 31 to 41 nm, confirming effective transmission of the applied nanovibrational stimulus through the overlying water.

An attempt was made to directly measure the motion of the zebrafish embryos (Figure 1G). LDV measurements acquired on embryos and adjacent background regions were comparable (Figure 1E), indicating that the vibrational signal detected at the embryo location was similar to that of the surrounding vibrating environment. Because the embryos were positioned close to the bottom surface, the recorded signal was dominated by motion of the underlying substrate, preventing direct isolation of embryo-specific displacement in this configuration. Nevertheless, these measurements are consistent with effective transmission of the nanovibrational stimulus to the embryo microenvironment.

### Nanovibration does not lead to gross morphological changes in zebrafish development

To determine whether nanovibrational stimulation affects overall development *in vivo*, zebrafish larvae were exposed to nanovibration from 3 to 5 days post-fertilisation (dpf). Gross morphological assessment revealed no overt developmental abnormalities in treated larvae compared with controls (Figure 2). Body and head length measurements at 5 dpf showed no significant difference between groups (Figure 2C), indicating that nanovibrational stimulation neither enhances nor impairs overall growth or gross developmental morphology during this period.

**Figure 2.**
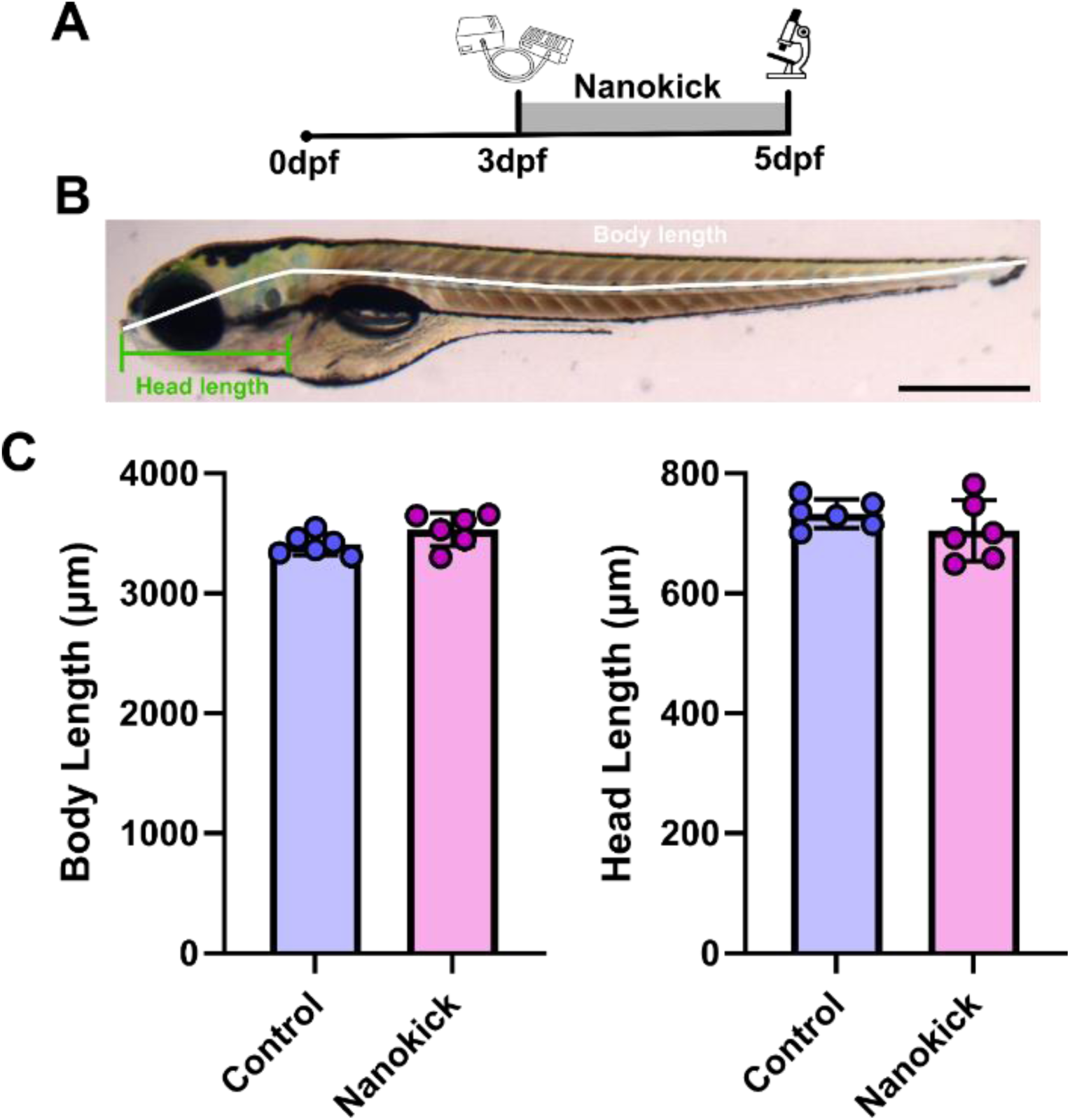
Exposure to nano-vibration does not lead to changes to body or head length. **(A)** Schematic depicting timecourse of larval nano-vibration treatment. **(B)** Representative lateral image of zebrafish showing body length measurement (white line) and head length (green line) at 5dpf. Scale bar =500µm. **(C)** Graphs showing measurement for physiological body length and head length from controls and nanovibration treated group at 5dpf. N = 6 per group

### Nanovibration enhances C2C12 muscle cell differentiation

Skeletal muscle is highly mechanosensitive. As the effects of nanovibration have been previously been characterized for muscle lineages. We assessed the differentiation of C2C12 cells in 2D culture following nanovibration treatment (Figure 3). Nanovibration-treated C2C12 cultures altered differentiation, with nanovibrated cells showing an approximately three-fold higher differentiation rate compared to non-stimulated cells, quantified as cells in myotubes, with Myosin heavy chain protein expression (Figure 3A-B), consistent with terminal myogenic differentiation.

**Figure 3.**
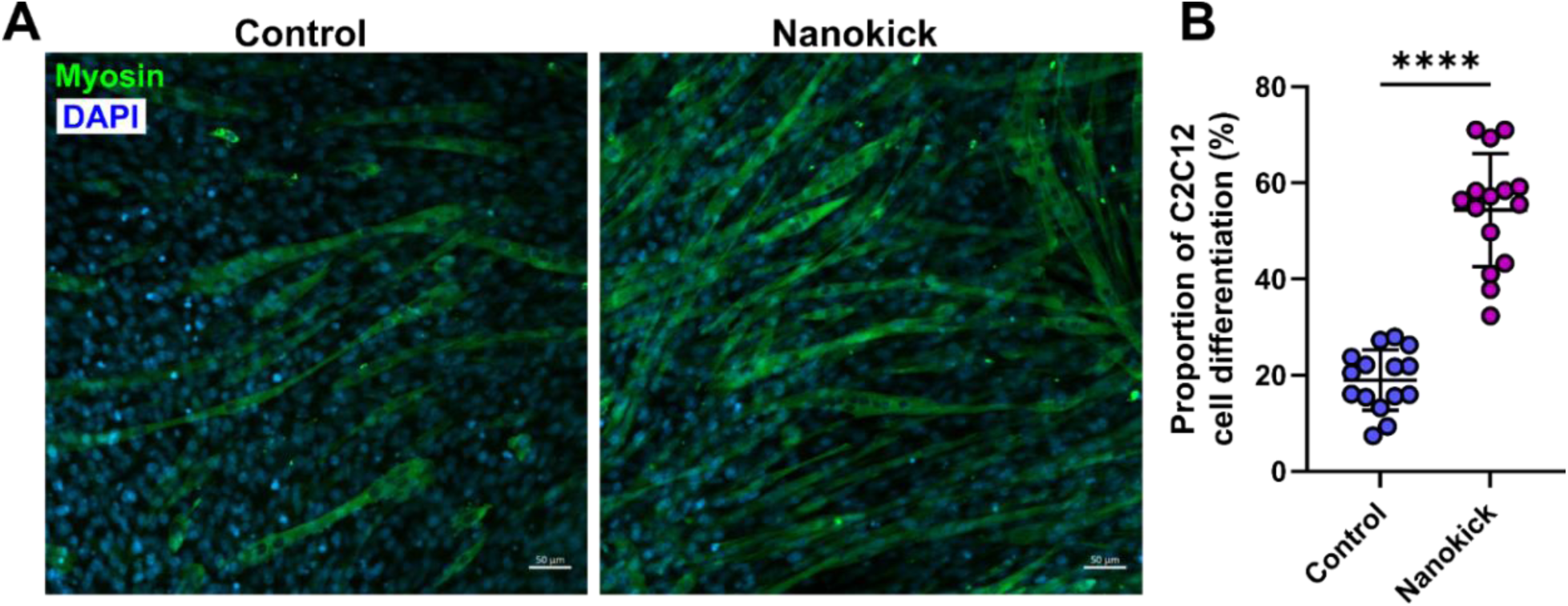
Nanovibration enhances C2C12 cell differentiation in 2D. **(A)** Representative confocal images ofC2C12 cells under control or nano-vibration treatment for 48 hours. Cells immunostained for myosin (green) and DAPI for nuclei (blue). Scale bars = 50µm.**(B)** Percentage of cell differentiation following control or nanovibration treatment. Welch’s T test performed where *\*\*\*\*P* <0.0001. N = 5 images per replicate and N= 3 replicates per group.

### Nanovibration does not significantly affect craniofacial or trunk muscle morphology in larval zebrafish

To test whether this effect was observed in vivo we quantified muscle after exposure of larvae to nanovibration from 3dpf to 5dpf (Figure 4). In craniofacial muscles, there was no significant change in muscle fibre length, width or number, or in muscle fibre proliferation in the intermandibularis (imp) or interhyal (ih) muscles between treated and control larvae (Figure 4C-G). Similarly, in the trunk musculature, no significant changes were observed in somite area or length or muscle fibre proliferation (Figure 4H-L), however, increased EdU incorporation was observed in the cells overlying the muscle (yellow arrowheads Fig 4K), suggestive of increases to proliferation in dermomytome-derived muscle progenitors^36–38^.

**Figure 4.**
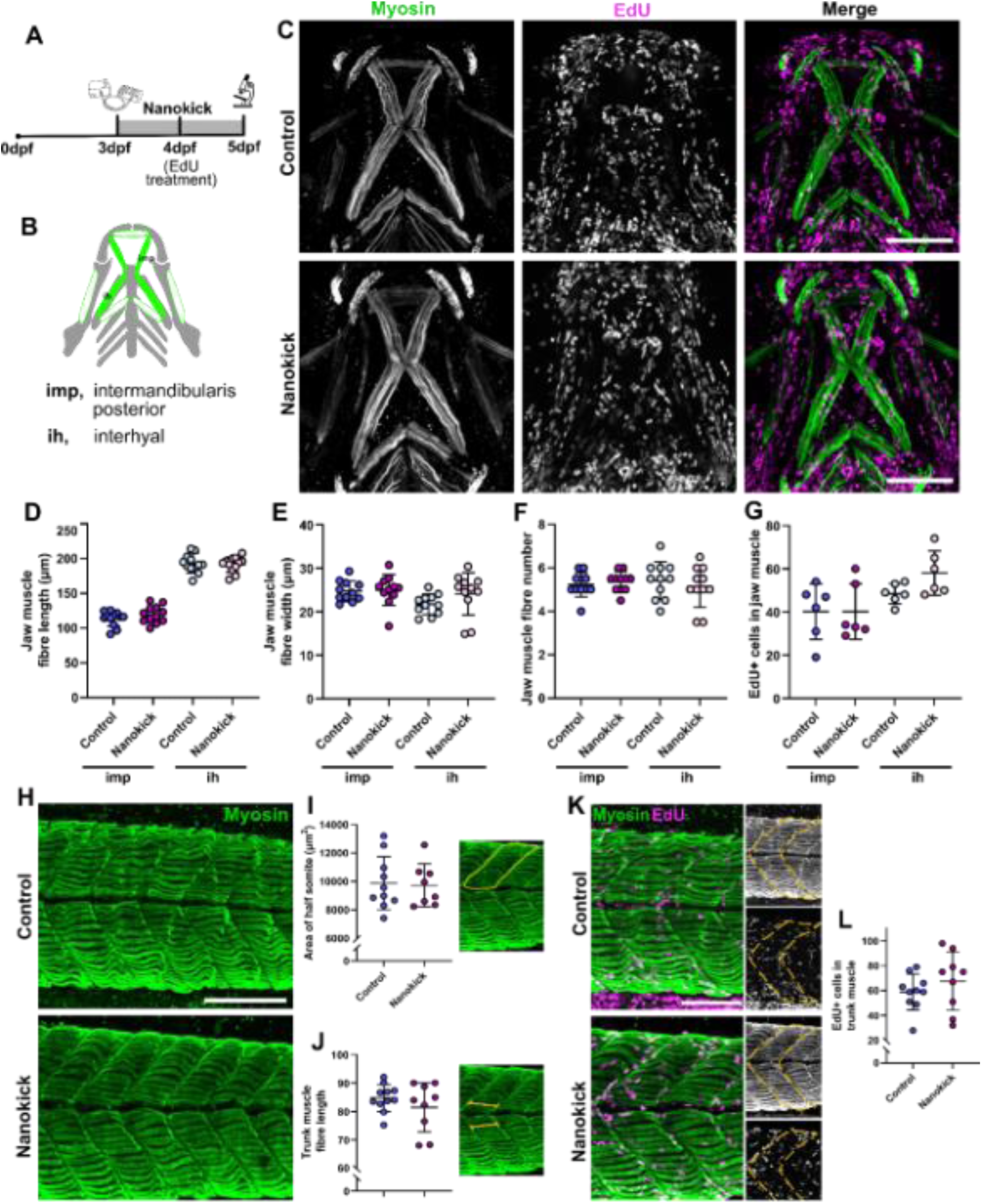
Craniofacial muscle formation of larvae is not affected by nano- vibration. **(A)** Schematic showing time course of nano-vibration treatment. **(B)** Schematic indicating main muscle groups measured (green) within the lower jaw (grey) at 5dpf. **(C)** Representative confocal images of lower jaw musculature in control and nanovibration-treated zebrafish immunostained for muscle (green) following 24-hour treatment with EdU Click-iT (magenta). Scale bars = 100µm.Quantification of **(D)** muscle fibre length, **(E)** muscle fibre width, **(F)** muscle fibre number and **(G)** number of EdU positive cells in the muscle fibres of IMP and IH muscles at 5dpf. N = 11per group for (D-F), N = 6 per group for (G). IMP, intermandibularis posterior; IH, interhyal. **(H)** Representative confocal max projections of trunk muscle in control and nanovibration-treated zebrafish immunostained for muscle (green)at 5dpf. Scale bars = 100µm. Quantification of **(I)** half somite area and **(J)** somite length. **(K)** Representative confocal max projections of trunk immunostained muscle (green) in control and nanokick-treated zebrafish at 5dpf following 24 hour treatment with EdU Click-iT (magenta). Yellow arrowheads mark putative dermomytome proliferation. Scale bars = 100µm. **(L)** Number of EdU positive cells in trunk muscle fibres (yellow dashed line shows region measured).

### Nanovibration in larval stages leads to increased bone deposition in the craniofacial skeleton

We next assessed whether nanovibration influences skeletal development by analysing operculum development (Figure 5). The operculum is one of the first dermal bones to become ossified in larvae and, therefore, can be used to assess early bone formation. Quantification of osx:eGFP-positive osteoblasts in the operculum revealed a significant increase in osteoblast number in nanovibration treated larvae compared with controls (Figure 5C-D). Consistent with this, overall operculum area and analysis of mineralised tissue demonstrated an increase in bone area within the operculum following nanovibrational stimulation, whilst operculum length remained unchanged (Figure 5E-G). These data indicate that nanovibration promotes osteogenic cell differentiation and bone formation in vivo without dramatic changes to bone morphology.

**Figure 5.**
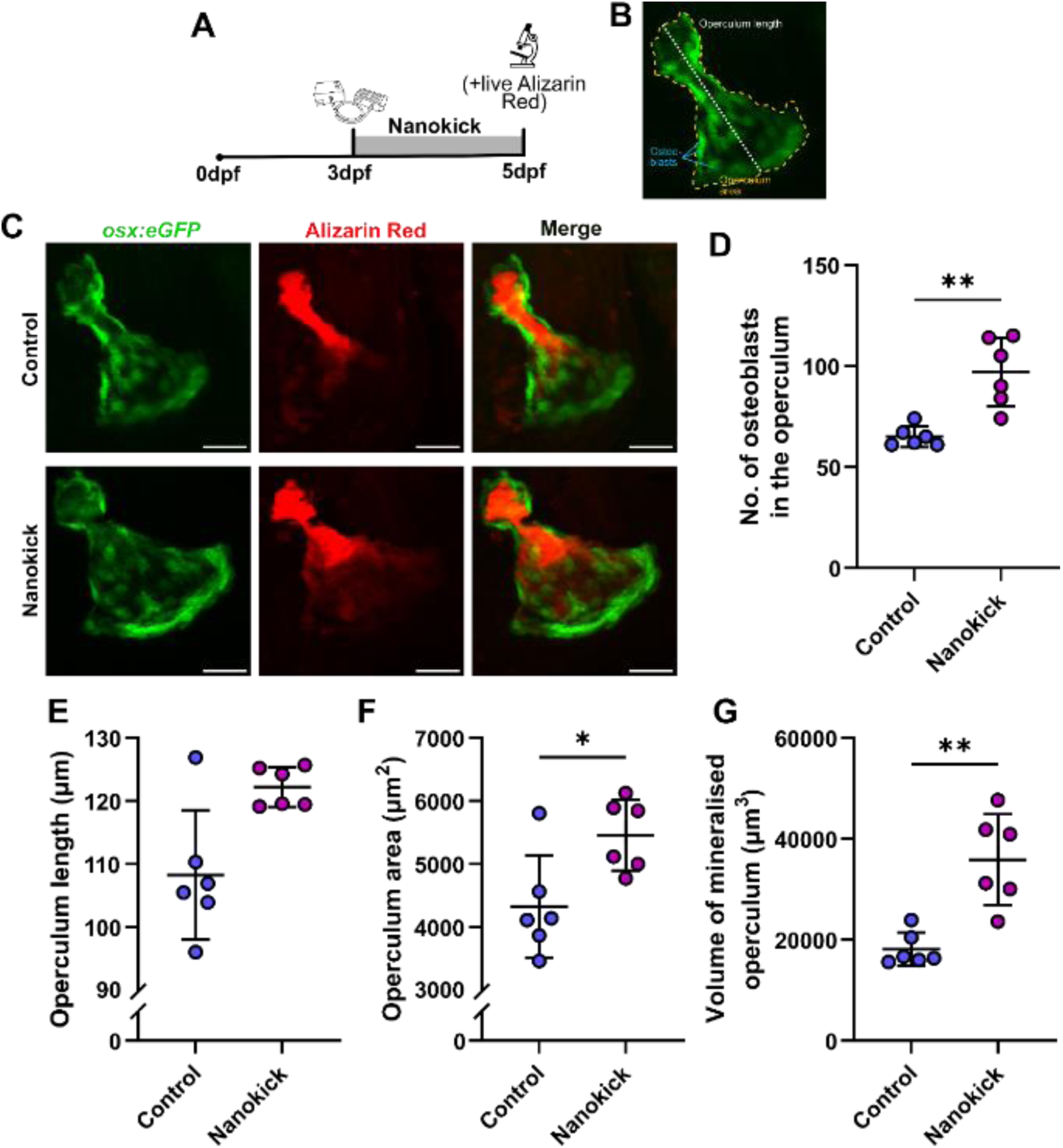
Nanovibration promotes osteoblast differentiation and bone mineralisation in larval operculum. **(A)** Schematic showing the time course of nano-vibration treatment. **(B)** Schematic showing operculum length and area measurements and identification of osteoblasts in the larval operculum at 5dpf. **(C)** Representative lateral images of the operculum in control and nanovibration treated *Tg(Ola. Sp7:nlsGFP)* zebrafish at 5dpf live stained with Alizarin red. Osteoblasts(green) and mineralised bone (alizarin red). Scale bars = 25µm.Graphs showing **(D)** number of osteoblasts in the operculum, **(E)** operculum length, **(F)** operculum area and **(G)** volume of mineralised bone in the operculum between control and nanovibration treated zebrafish at 5dpf. Welch’s t test performed for(D-F) where *\*\*P=*.0045 and *\*P=*.0204. Mann-Whitney test performed for (G) where \*\**P=*.0043. N = 6 for all.

### Nanovibration subtly affects craniofacial cartilage morphology

To determine whether nanovibration broadly affects all skeletal tissues, we examined craniofacial cartilage development. Cartilage morphology appeared broadly comparable between nanovibrated and control larvae (Figure 6A-D). However, there was a significant change at the joint site, with nanovibrated larvae showing increased joint neck width but decreased joint spacing (Figure 6E-G). Concomitantly, there was a minor increase in proliferation at the joint site, suggesting a subtle effect on proliferation and organisation of chondrocytes at the joint site, which is a region of high strain across the 3-5 dpf period due to onset of jaw movement ^39,40^.

**Figure 6.**
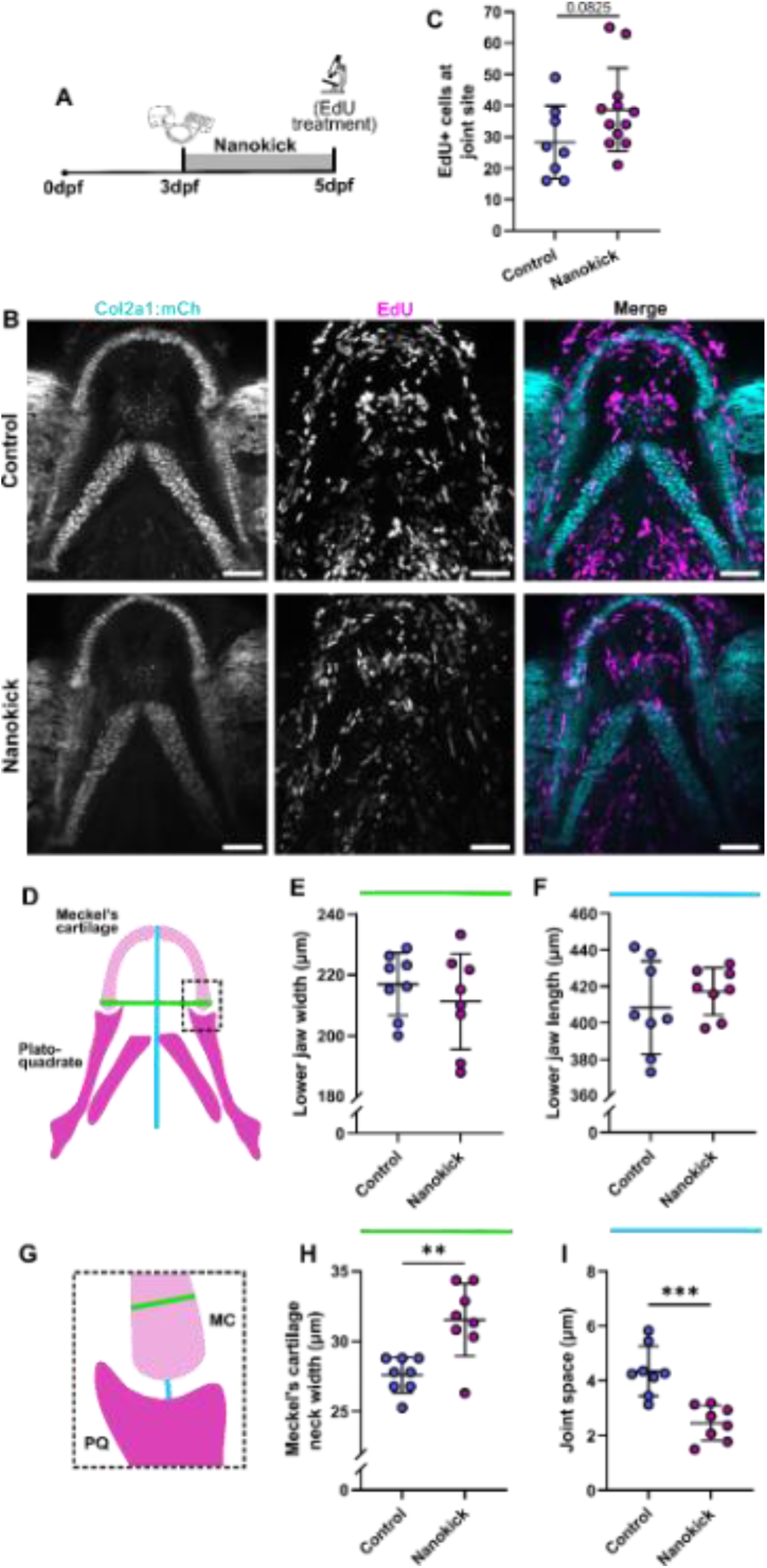
Nano-vibration alters lower jaw and joint morphology in larval development. **(A)** Schematic showing time course of nanovibration treatment. **(B)** Representative confocal images of z-stack projections of the lower jaw at 5dpfin control and nanovibration-treated *Tg(Col2a1:mCherry)* larvae(cyan) treated with EdU Click-iT (magenta) for 24 hours from 4-5dpf. Scale bars = 50μm. **(C)** Quantification of number of EdU-positive cells at the joint site. N = 8 for control and 12 for nanovibration treated. Welch’s T test performed. **(D)** Schematic of zebrafish lower jaw showing lower jaw length (blue) and width (green)measurements. Quantification of **(C)** lower jaw width and **(E)** length. **(F)** Inset from black dotted box in (B) showing jaw joint width (green) and joint space (blue) measurements. MC, Meckel’s cartilage; PQ, palatoquadrate. Quantification of **(G)** Meckel’s cartilage neck width and **(G)** joint space. Welch’s T test performed for both where *\*\*P=.*0031 and *\*\*\*P=*.0004.

### Nanovibration does not lead to changes in basal immune cell number or localization

As there are known interactions between osteogenic and immune cell lineages, we wanted to establish whether nanovibration leads to changes in basal innate immune cell number. We quantified mpx:eGFP-positive neutrophils and mpeg:mCherry-positive macrophages in the head region of larvae following exposure to nanovibration from 3–5 dpf (Figure 7). Overall, nanovibration did not alter the gross spatial localisation of innate immune cells (Figure 7A,B). A small decrease in neutrophil number was observed at 48 hours, while macrophage number was not significantly altered (Figure 7C-D). These data suggest that nanovibration does not induce ectopic innate immune cell accumulation or a basal inflammatory response under these conditions.

**Figure 7.**
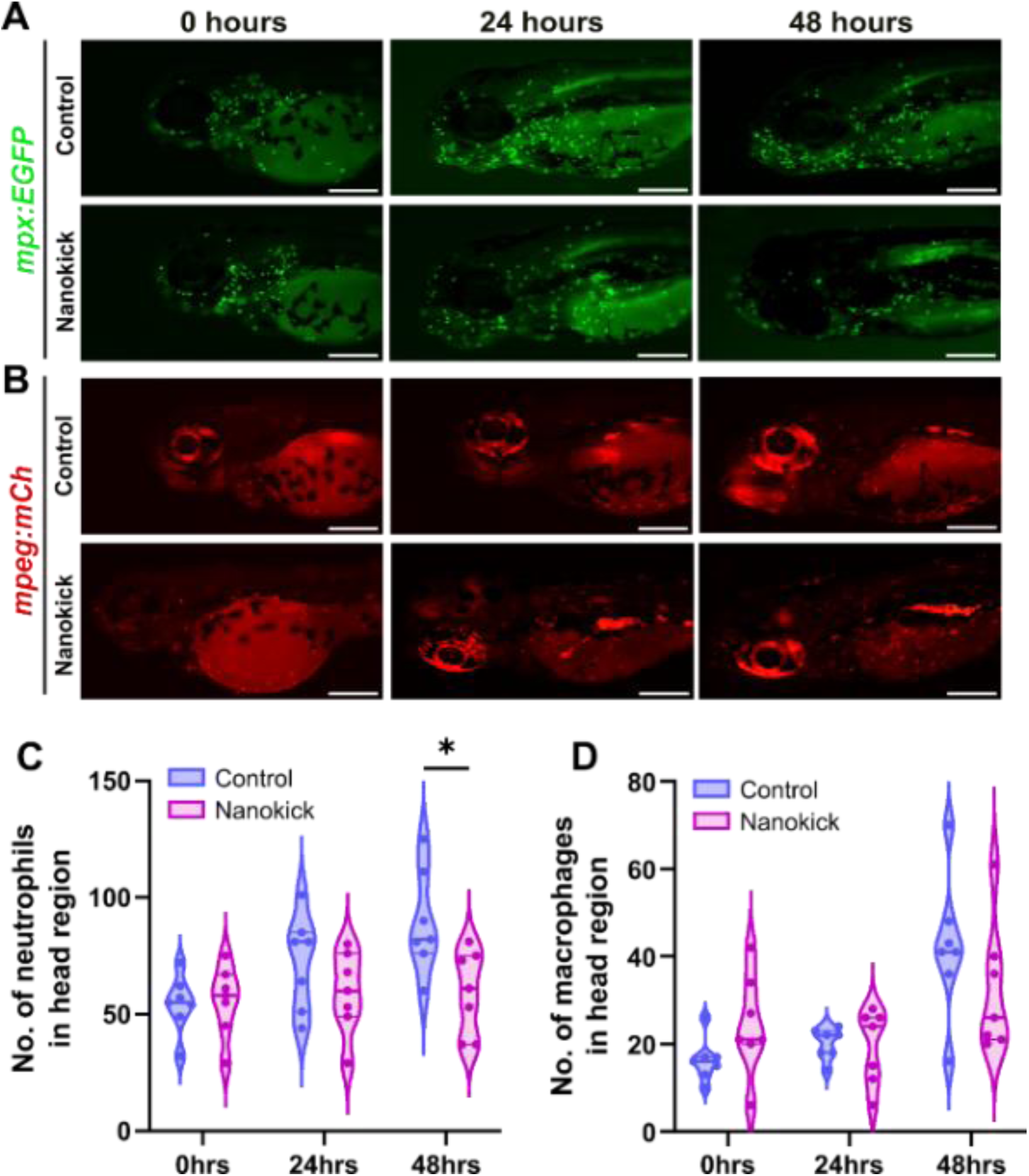
Basal immune cell levels are unaffected by nanovibration treatment. Representative stereomicroscope images of. **(A)** *TgBAC(mpx:GFP)* and **(B)** *Tg(mpeg1:mCherry)* positive control and nanovibration-treated larvae at 3-5dpf. Scale bars = 200µm. **(C)** Neutrophils (green) and **(D)** macrophages (red) were counted in the head region at 0, 24 and 48 hours post control or nanovibration treatment. Two-way ANOVA performed for both where *\*P=.0106.* N = 7 for all groups and timepoints.

### Nanovibration leads to increased osteoblast number following bone injury in scales

Having established that nanovibration preferentially activated osteogenesis in a larval model, we wanted to test whether these pro-osteogenic cellular effects were preserved in an adult context. We therefore established an *ex vivo* scale injury model in which scales from transgenic fish carrying the sp7 reporter were harvested, injuries were made with a scalpel then the whole scale was cultured for 48 hours with or without nanovibration (Figure 8A,B). Nanovibration-treated scales showed significantly increased sp7:eGFP mean pixel intensity at the injury site compared with controls (Figure 8C). These data suggest that nanovibration increases osteoblast reporter activity at adult scale injury sites, consistent with an enhanced osteogenic repair response.

**Figure 8.**
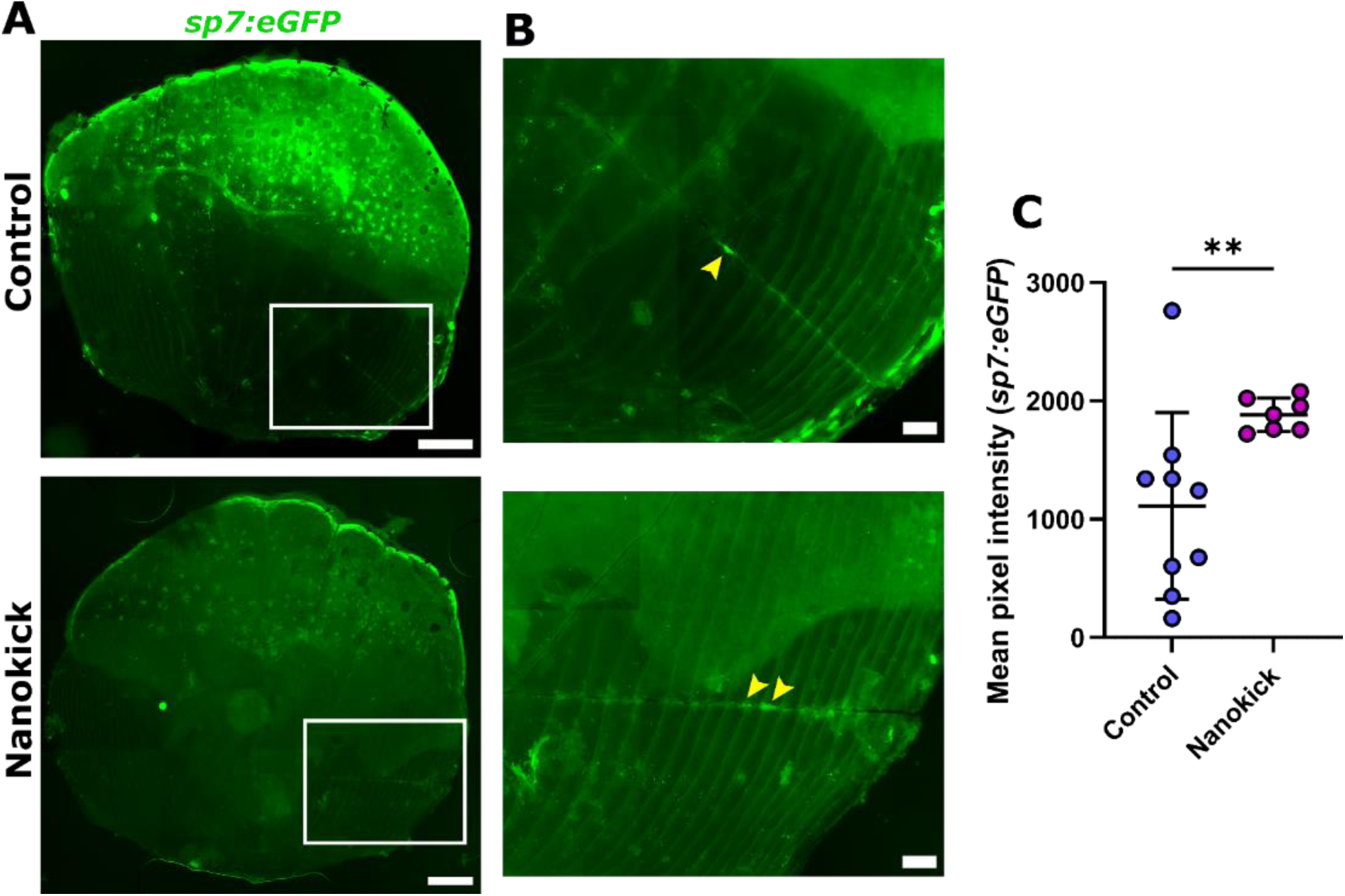
Nanovibration promotes osteoblast expression at fracture sites in scales. **(A)** Representative slide scanner images of scales from *Tg(Ola. Sp7:nlsGFP)* fish at 10mpf. Scales were fractured *ex vivo* and cultured for 48 hours under control or nanovibration conditions prior to imaging. White box shows location of fracture site as shown in **(B).** Yellow arrowheads show osteoblasts at injury site. Scale bars = 200µm and 50µm, respectively. **(C)** Mean pixel intensity of osteoblast fluorescence at fracture site. Mann-Whitney test performed where \*\**P=*.0079. Six scales from 3 fish each were taken and split evenly between treatment groups (N = 9scales per group). Two measurements from nanovibration treated group were removed as outliers.

## Discussion

In this study, we demonstrate that nanovibrational stimulation enhances osteogenesis *in vivo* in zebrafish, as evidenced by increased osteoblast number and bone formation in larvae and increased osterix expression at a bone injury site in *ex vivo* culture. Importantly, these effects occur in the absence of substantial changes in other tissues, including skeletal muscle, craniofacial cartilage, and innate immune cell populations. Together, these findings support the hypothesis that continuous sinusoidal nanovibration with a amplitude of 30 nm, and frequency of 1 kHz acts as a selective osteogenic mechanobiological cue, promoting osteogenic processes while exerting minimal broader physiological disruption. Our results extend previous *in vitro* observations by showing that nanovibrational stimulation can drive osteogenic outcomes within a whole-organism context resulting as increased osteoblast activity and enhanced mineralisation.

We show for the first time that nanovibration can promote myogenic differentiation *in vitro*. However, a key finding of this study is the preferential nature of the osteogenic response *in vivo*. Despite clear effects on bone formation, we observed limited changes to other cellular systems, including skeletal muscle, where effects in in vivo were minimal. This suggests that nanovibration does not act as a generalised growth or stress stimulus, but rather stimulates cells in some tissues more readily than others, at least at the developmental stages studied.

One explanation may lie in the variable mechanosensitivity of different cell types, but could also reflect differences in cellular context and the microenvironment. The sensitivity of cells to mechanical stimulation is known to differ between 2D and 3D environments ^41^. The differences between the in vitro response of C2C12 cells in culture and skeletal muscle in vivo could reflect changes to microenvironment; muscle differences to proliferation have been observed upon changes to matrix stiffness^42^. Alternatively, it could reflect the timing that the stimulus was applied. In our culture experiments the cells were confluent but not yet differentiated. Whereas trunk muscle in zebrafish forms early in development with trunk muscle fibres formed and innervated prior to application of nanovibrational stimulus^43,44,45^. Even within the jaw, where muscle forms later, functional muscles are present prior to application of the nanovibration in our experiments^40 46^. Together these observations suggest cellular responsivenss may dependent not only on cell type but also on developmental state and tissue context.

A similar stage dependent response may explain underpin the relatively subtle effects seen on craniofacial cartilage. Significant changes were restricted to the jaw joint regionwith more mature chondrocytes less affected. Cells in the jaw joint alter orientation in response to changes to muscle-activity during this developmental window (3-5 dpf), consistent with these chondrocytes being chondrocytes being the most mechanosensitive in this period^25^. Mechanical strain, generated by muscular force, is well established as a regulator of skeletal development and ossification across vertebrates, with loss of strain leading to reduced ossification and altered growth plate progression^3,4,48^. These findings raise the possibility that nanovibration engages mechanosensitive pathways that overlap with those activated by endogenous mechanical stimuli.

Although the present study does not directly interrogate underlying mechanisms, our findings are consistent with emerging models of nanovibrational mechanotransduction. In vitro work have shown nanovibration induces cytoskeletal reorganisation, increases intracellular tension, and activates mechanosensitive signalling pathways, including calcium flux and downstream transcriptional regulators of osteogenesis^5,49^. Recent work has also implicated metabolic reprogramming and reactive oxygen species (ROS) signalling as contributors to nanovibration-induced osteogenesis^6,18^. Within an *in vivo* context, these mechanisms may enhance osteoblast differentiation and activity without triggering widespread stress or inflammatory responses. The absence of major changes in immune cell number in our study is consistent with a model in which nanovibration operates within a physiological rather than pathological range of mechanical stimulation. Future work examining gene expression, ion channel activity, and metabolic state *in vivo* will be important to elucidating these mechanisms.

In summary, we show that nanovibrational stimulation enhances osteogenesis in zebrafish in vivo while exerting minimal effects on muscle, cartilage and immune systems at the developmental stages studied. These findings establish that nanoscale mechanical stimulation can promote bone formation within a living vertebrate system and support the further development of nanovibration as a non-invasive strategy for enhancing skeletal repair. More broadly, our results demonstrate that subtle mechanical cues can elicit distinct tissue-specific responses in vivo, providing new insight into how different cell populations interpret and respond to mechanical information during development and repair.

## Acknowledgements

We thank the staff of the Wolfson Bioimaging Facility for imaging support and the MRC grant (MRC MC_PC_MR/X01391X/1) for funding the Evident VS200 slide scanner. We also thank Mathew Green and technical staff from the University of Bristol’s Animal Scientific Unit (ASU) for providing zebrafish husbandry. We also thank Prof James Windmill for access to vibrometry facilities.

## Author contributions

J.A. Williams and C.L. Hammond conceptualised the study and designed experiments. O. Adigun, M. Wang, J. Zappia, M. Maung, J.A Williams, O. Dobre and J.J. Moss conducted the experiments and interpreted data. P.G. Childs and S. Reid contributed resources (nanovibration devices) for the work. O. Adigun, M. Wang, J.A. Williams and J.J. Moss performed statistical analysis and made all figures. C.L. Hammond and J.J. Moss wrote the first draft of the manuscript, and all authors made intellectual contributions and assisted in manuscript editing.

## Data availability

All raw data will be available via a DOI at data.bris.ac.uk upon article acceptance.

## References

1. Lacroix, D. & Prendergast, P. J. A mechano-regulation model for tissue differentiation during fracture healing: Analysis of gap size and loading. J. Biomech. 35, 1163–1171 (2002).

2. Boerckel, J. D. et al. Effects of in vivo mechanical loading on large bone defect regeneration. J. Orthop. Res. (2012) doi:10.1002/jor.22042.

3. Nowlan, N. C., Sharpe, J., Roddy, K. A., Prendergast, P. J. & Murphy, P. Mechanobiology of embryonic skeletal development: Insights from animal models. Birth Defects Res C Embryo Today 90, 203–213 (2010).

4. Nowlan, N. C., Murphy, P. & Prendergast, P. J. Mechanobiology of embryonic limb development. Ann N Y Acad Sci 1101, 389–411 (2007).

5. Pemberton, G. D. et al. Nanoscale stimulation of osteoblastogenesis from mesenchymal stem cells: Nanotopography and nanokicking. Nanomedicine 10, 547–560 (2015).

6. Orapiriyakul, W. et al. Nanovibrational Stimulation of Mesenchymal Stem Cells Induces Therapeutic Reactive Oxygen Species and Inflammation for Three-Dimensional Bone Tissue Engineering. ACS Nano 14, 10027–10044 (2020).

7. Williams, J. A. et al. Developing and Investigating a Nanovibration Intervention for the Prevention/Reversal of Bone Loss Following Spinal Cord Injury. ACS Nano 18, 17630–17641 (2024).

8. Cardinale, M. & Rittweger, J. Vibration exercise makes your muscles and bones stronger: fact or fiction? J. Br. Menopause Soc. 12, 12–18 (2006).

9. Lin, Y. H. et al. The essential role of stathmin in myoblast c2c12 for vertical vibration- induced myotube formation. Biomolecules 11, (2021).

10. Carrasco, V. B., Vidal, J. M. & Caparrós-Manosalva, C. Vibration motor stimulation device in smart leggings that promotes motor performance in older people. Med. Biol. Eng. Comput. 61, 635–649 (2023).

11. Sato, S. et al. Vibration acceleration enhances proliferation, migration, and maturation of C2C12 cells and promotes regeneration of muscle injury in male rats. Physiol. Rep. 12, (2024).

12. Mittermayr, R., Haffner, N., Feichtinger, X. & Schaden, W. The role of shockwaves in the enhancement of bone repair - from basic principles to clinical application. Injury 52 **Suppl 2**, S84–S90 (2021).

13. Padilla, F., Puts, R., Vico, L. & Raum, K. Stimulation of bone repair with ultrasound: a review of the possible mechanic effects. Ultrasonics 54, 1125–1145 (2014).

14. Rubin, C. T. et al. Prevention of postmenopausal bone loss by a low-magnitude, high-frequency mechanical stimuli: a clinical trial assessing compliance, efficacy, and safety. J. Bone Miner. Res. 19, 343–351 (2004).

15. Campsie, P. et al. Design, construction and characterisation of a novel nanovibrational bioreactor and cultureware for osteogenesis. Sci. Rep. 9, (2019).

16. Yao, M. et al. Mechanical activation of vinculin binding to talin locks talin in an unfolded conformation. Sci. Rep. 4, (2014).

17. Kennedy, J. W. et al. Nanovibrational stimulation inhibits osteoclastogenesis and enhances osteogenesis in co-cultures. Sci. Rep. 11, (2021).

18. Hodgkinson, T. et al. The use of nanovibration to discover specific and potent bioactive metabolites that stimulate osteogenic differentiation in mesenchymal stem cells. Sci. Adv. 7, (2021).

19. Bispo, D. S. C. et al. Global metabolomics identifies new extracellular biomarkers of nanovibration-driven mesenchymal stem cells osteodifferentiation. Biomater. Adv. 180, (2026).

20. Orapiriyakul, W. et al. Nanovibrational Stimulation of Mesenchymal Stem Cells Induces Therapeutic Reactive Oxygen Species and Inflammation for Three-Dimensional Bone Tissue Engineering. ACS Nano 14, 10027–10044 (2020).

21. Dietrich, K. et al. Skeletal Biology and Disease Modeling in Zebrafish. J. Bone Miner. Res. 36, 436–458 (2021).

22. Tonelli, F. et al. Zebrafish: A Resourceful Vertebrate Model to Investigate Skeletal Disorders. Front. Endocrinol. (Lausanne). (2020) doi:10.3389/fendo.2020.00489.

23. Brunt, L. H., Begg, K., Kague, E., Cross, S. & Hammond, C. L. Wnt signalling controls the response to mechanical loading during zebrafish joint development. Dev. 144, (2017).

24. Brunt, L. H., Roddy, K. A., Rayfield, E. J. & Hammond, C. L. Building finite element models to investigate zebrafish jaw biomechanics. J. Vis. Exp. 2016, (2016).

25. Brunt, L. H. et al. Differential effects of altered patterns of movement and strain on joint cell behaviour and skeletal morphogenesis. Osteoarthr. Cartil. 24, 1940–1950 (2016).

26. Tong, Q., Raele, R., Bergen, D. & Hammond, C. Zebrafish Scale Regeneration In Toto and Ex Vivo Culture of Scales. J. Vis. Exp. 2024, (2024).

27. Bergen, D. J. M. et al. Regenerating zebrafish scales express a subset of evolutionary conserved genes involved in human skeletal disease. BMC Biol. 20, (2022).

28. Bergen, D. J. M., Kague, E. & Hammond, C. L. Zebrafish as an emerging model for osteoporosis: A primary testing platform for screening new osteo-active compounds. Front. Endocrinol. (Lausanne). 10, (2019).

29. De Vrieze, E., Moren, M., Metz, J. R., Flik, G. & Lie, K. K. Arachidonic acid enhances turnover of the dermal skeleton: Studies on zebrafish scales e89347. PLoS One 9, (2014).

30. Knopf, F. et al. Bone regenerates via dedifferentiation of osteoblasts in the zebrafish fin. Dev. Cell (2011) doi:10.1016/j.devcel.2011.04.014.

31. Mitchell, R. E. et al. New tools for studying osteoarthritis genetics in zebrafish. Osteoarthr. Cartil. 21, 269–278 (2013).

32. von Hofsten, J. et al. Prdm1- and Sox6-mediated transcriptional repression specifies muscle fibre type in the zebrafish embryo. EMBO Rep 9, 683–689 (2008).

33. Kimmel, C. B., Sessions, S. K. & MacLeod, M. C. Evidence for an association of most nuclear RNA with chromatin. J Mol Biol 102, 177–191 (1976).

34. Renshaw, S. A. et al. A transgenic zebrafish model of neutrophilic inflammation. Blood 108, 3976–3978 (2006).

35. Schindelin, J., et al. Fiji: an open-source platform for biological-image analysis. Nat. Methods 9, 676–82 (2012).

36. Roy, S. D. et al. Myotome adaptability confers developmental robustness to somitic myogenesis in response to fibre number alteration. Dev. Biol. 431, (2017).

37. Devoto, S. H. et al. Generality of vertebrate developmental patterns: Evidence for a dermomyotome in fish. Evol. Dev. 8, (2006).

38. Hammond, C. L. et al. Signals and myogenic regulatory factors restrict pax3 and pax7 expression to dermomyotome-like tissue in zebrafish. Dev. Biol. 302, (2007).

39. Brunt, L. H., Norton, J. L., Bright, J. A., Rayfield, E. J. & Hammond, C. L. Finite element modelling predicts changes in joint shape and cell behaviour due to loss of muscle strain in jaw development. J. Biomech. 48, 3112–3122 (2015).

40. Brunt, L. H. et al. Differential effects of altered patterns of movement and strain on joint cell behaviour and skeletal morphogenesis. Osteoarthr. Cartil. 24, (2016).

41. Saraswathibhatla, A., Indana, D. & Chaudhuri, O. Cell-extracellular matrix mechanotransduction in 3D. Nat. Rev. Mol. Cell Biol. 24, 495–516 (2023).

42. Trensz, F. et al. Increased microenvironment stiffness in damaged myofibers promotes myogenic progenitor cell proliferation. Skelet. Muscle 5, 5 (2015).

43. Blagden, C. S., Currie, P. D., Ingham, P. W. & Hughes, S. M. Notochord induction of zebrafish slow muscle mediated by Sonic hedgehog. Genes Dev 11, 2163–2175 (1997).

44. Groves, J. A., Hammond, C. L. & Hughes, S. M. Fgf8 drives myogenic progression of a novel lateral fast muscle fibre population in zebrafish. Development 132, (2005).

45. Kimmel, C. B., Hatta, K. & Eisen, J. S. Genetic control of primary neuronal development in zebrafish. Development **Suppl** 2, 47–57 (1991).

46. Schilling, T. F. & Kimmel, C. B. Musculoskeletal patterning in the pharyngeal segments of the zebrafish embryo. Development 124, 2945–2960 (1997).

47. Pollard, A. S., McGonnell, I. M. & Pitsillides, A. A. Mechanoadaptation of developing limbs: shaking a leg. J Anat 224, 615–623 (2014).

48. Roddy, K. A., Kelly, G. M., van Es, M. H., Murphy, P. & Prendergast, P. J. Dynamic patterns of mechanical stimulation co-localise with growth and cell proliferation during morphogenesis in the avian embryonic knee joint. J Biomech 44, 143–149 (2011).

49. Pereira Sousa, R. P., Reid, S., Dalby, M. J., Jimenez, M. & Childs, P. G. Mechanical Phenotyping of MG63s Following Vibrational Stimulation. FASEB J. 40, e71807 (2026).

